# Sleep loss overrides sleep-preparatory behavior to promote sleep

**DOI:** 10.1101/2025.06.02.657276

**Authors:** Nicole Freitag, Kareem El-Tawil, Aaron Crane, Vanessa Hilo, George Saad, Ada Eban-Rothschild

**Affiliations:** Department of Psychology, University of Michigan, Ann Arbor, MI 48109, USA

**Keywords:** Pre-sleep behavior, nest-building, sleep deprivation, de-arousal, stress

## Abstract

In the period preceding sleep, humans and other animals display a stereotyped repertoire of behaviors–including hygiene-related activities and preparing a place to sleep. Evidence suggests that this pre-sleep phase actively contributes to sleep initiation and quality. Nonetheless, individuals can sometimes fall asleep without preparation, even under undesirable circumstances. These abrupt transitions into sleep can have severe consequences, particularly in high-risk environments. Although progress has been made in identifying neuronal populations controlling sleep/wake states and mechanisms regulating cortical oscillations during sleep, little is known about the natural processes that govern the pre-sleep phase, under baseline conditions and following sleep loss. Here, we examine factors regulating pre-sleep behaviors using environmental and behavioral manipulations, video recordings, machine-learning-based tracking, and EEG-EMG analysis in freely-behaving mice. We focus on nest-building–a key pre-sleep behavior–and assess its modulation by time of day and sleep deprivation. We find that mice are highly motivated to build nests during the light phase but show reduced motivation during most of the dark phase. Sleep deprivation significantly suppresses pre-sleep nest-building and promotes the direct initiation of sleep. Varying amounts of sleep deprivation, from 2–6 hours, uniformly suppress nest-building. This suppression is not due to stress, as mice exposed to acute restraint stress engage robustly in nest-building. Our findings provide insight into processes regulating the transition from wakefulness to sleep. Understanding pre-sleep regulation has important implications for treating sleep-onset difficulties–prevalent in insomnia and predictive of cognitive decline–and for mitigating risks associated with uncontrolled sleep onset in high-stakes situations.

**STATEMENT OF SIGNIFICANCE:** Prior to sleep, animals–including humans–engage in stereotyped behaviors such as preparing a sleep space, thought to facilitate sleep initiation and quality. However, sleep can occur abruptly without preparation, posing serious risks, especially in high-stakes environments. Despite progress in understanding sleep/wake circuitry, the regulation of pre-sleep behaviors remains poorly understood. Here, we used multiple techniques in mice to examine nest-building, a key pre-sleep behavior. We found that mice are more motivated to build nests during the light phase, but this behavior is strongly suppressed by sleep loss and is not affected by acute stress. These findings highlight pre-sleep behaviors as an active, regulated component of sleep onset, with implications for understanding and treating sleep initiation difficulties.

## INTRODUCTION

In the period immediately preceding the sleep phase, humans and other animals display a stereotypical repertoire of behaviors, including locating and preparing a safe sleeping area, performing hygiene-related behaviors, and assuming a sleeping posture^1–5^. The pre-sleep period is recognized as a transitional phase that promotes relaxation and actively contributes to the initiation and continuity of sleep^6–12^. Nonetheless, individuals can sometimes fall asleep without any preparation, even under entirely undesirable circumstances, such as while driving. These abrupt transitions into sleep, typically attributed to sleepiness (e.g., ^13^), can have severe consequences for both the individuals involved and society when they occur in risk-prone environments. Although significant efforts have been made to elucidate the brain mechanisms that regulate cortical oscillatory activity during sleep and to pinpoint neuronal populations controlling sleep/wake states^7,14,15^, surprisingly little is known about the natural processes that govern the pre-sleep phase and the initiation of sleep both under baseline conditions and following sleep loss.

Sleep timing, depth, and duration are understood to be regulated by two key processes: the circadian clock and the homeostatic sleep drive^16,17^. The circadian process generates a 24-hour cycle in physiology and behavior, setting the sleep/wake schedule^17–19^. In contrast, the homeostatic process is thought to gauge sleep need based on previous durations of wakefulness, with longer wake periods leading to increased sleep need^17,19,20^. While much effort has been directed toward elucidating the functions of circadian and homeostatic processes in the regulation of sleep timing, duration and depth, the impact of these processes on pre-sleep behaviors remains unexplored.

Our study aims to fill this knowledge gap by characterizing the factors that control behavioral preparation for sleep. To achieve this, we utilized environmental and behavioral manipulations, video recording, machine-learning-based tracking, and EEG-EMG analysis in freely behaving and sleeping mice. We determined the effect of time of day and sleep deprivation on the motivation to build a nest–a key pre-sleep behavior in mice^6,21^–as well as on locomotion and sleep/wake architecture. Motivation refers to internal processes that initiate and sustain goal-directed behavior. We inferred the motivation for nest-building from nest quality, which reflects the intensity and duration of nesting behavior. We found that mice exhibit a strong motivation to build nests throughout the light phase, while showing little motivation at the beginning of the dark phase. Additionally, we found that sleep loss suppresses nest-building behavior while also reducing locomotion and accelerating sleep initiation. Moreover, varying amounts of sleep deprivation–from 2 to 6 hours–uniformly suppress nest-building, suggesting that the mechanisms responsible for this behavioral suppression do not act proportionally to the amount of sleep lost, in contrast to the proportional nature of the homeostatic sleep rebound. Notably, stress alone does not account for this suppression of nesting behavior, as evidenced by the strong motivation to build a nest following acute restraint stress. Our findings significantly enhance current understanding of the processes regulating the transition from wakefulness to sleep. A better understanding of these phenomena is crucial, not only because abrupt and uncontrolled transitions into sleep can have severe consequences for both individuals and society, but also because difficulties in initiating sleep are prominent symptoms of insomnia^24^ and strong predictors of cognitive decline later in life^23,24^.

## METHODS

### Animals

We used in-house bred wild-type (WT) C57BL/6J mice (The Jackson Laboratory, Stock #: 000664) that were reproductively inexperienced and at least 8 weeks old. The mice were maintained under a 12-hr light/dark cycle at 22 ± 1°C, and provided with compressed cotton ‘Nestlet’ material (Ancare, Bellmore, NY, U.S.A.) and ad libitum access to water and food. Throughout the experiments, mice were individually housed in custom Plexiglas recording chambers (28.6 (W) × 39.4 (L) × 19.3 (H) cm) mounted with USB cameras (1080p USB webcam, Angetube). All experimental procedures adhered to the US National Institutes of Health Guide for the Care and Use of Laboratory Animals and were approved by the University of Michigan’s Institutional Animal Care and Use Committee.

### EEG-EMG implantation

Mice were anesthetized using a ketamine-xylazine mixture (100 and 10 mg kg^-1^, respectively) delivered via intraperitoneal injection (IP). Following this, they were administered lidocaine and carprofen (4 mg kg^-1^ and 5 mg kg^-1^, respectively). Subsequently, the mice were positioned in a stereotaxic frame (David Kopf Instruments, Tujunga, CA, U.S.A.) and maintained under anesthesia with approximately 1% isoflurane in O_2_. Mice were fitted with two miniature screw electrodes (F00CE125, J.I. Morris miniature fasteners, Inc.) at coordinates AP = 1.6 mm, ML = -

1.6 mm and AP = -2.5 mm, ML = -2.8 mm, and with two EMG wire electrodes (A S633, Cooner Wire) previously soldered to a 4-pin connector (custom). EMG electrodes were inserted between the trapezius muscles of the mice. The implant was anchored securely to the skull using C&B Metabond (Parkell) and dental cement, and the skin incision was closed using surgical sutures. After the surgery, the mice were placed on a heating pad until they regained full mobility. They were then given 7–10 days for recovery, after which they were transferred to individual open-top recording chambers and habituated to flexible EEG-EMG cables for an additional 10 days before data collection began.

### Nest manipulations and scoring

Manipulations were performed on mice that had resided in their home cages for at least three days. On experimental days, we removed the nests from the home cages of test mice and introduced new nesting material, consisting of four Nestlet strips weighing a total of 5 g. The new nesting material was placed away from the original nest location. Throughout the nest manipulation experiment, we recorded the mice using USB cameras operated with the iSpy software (iSpyConnect.com). Repeated nest manipulations involving the same mice were conducted at least four days apart.

Two hours after the introduction of the new nesting material, we visually inspected the nests and photographed the cages from the top and the side. We also weighed any unshredded Nestlet material. Nest quality was assessed using a six-point scale, which was modified from the scale presented in Deacon, 2006^27^:

1. Less than 5% of the Nestlet strips are shredded. The strips are typically dispersed across the home cage or left undisturbed in the original location where they were placed.
2. Between 5-15% of the Nestlet strips are shredded. The strips may be dispersed or gathered.
3. Between 15-30% of the Nestlet strips are shredded. The material is gathered within a quarter of the cage floor area, forming a discernible but flat nest.
4. 30-60% of the Nestlet strips are shredded and gathered to form either a platform or a cup-like structure. If a cup nest formation is present, the shredded material may be as little as 20%.
5. 60-80% of the Nestlet strips are torn and gathered. The walls of the nest are taller than the height of a mouse curled up on its side but cover less than 50% of the nest’s circumference.
6. More than 80% of the Nestlet strips are shredded. The nest resembles a crater with walls higher than a curled-up mouse, covering over 50% of its circumference.

Nest-scoring data from one female mouse were excluded because the animal failed to build a nest (score = 1) under any condition, including baseline.

### Sleep deprivation through environmental enrichment

Mice were placed in an enclosure containing bedding, water, and a running wheel, but no nesting material or food. They were continuously monitored by an experimenter in an adjacent room using the iSpy software (iSpyConnect.com) and, when available, their EEG-EMG signals. Whenever a mouse showed signs of quiescence, novel objects (e.g., wooden cubes, plastic figures) were introduced or exchanged in the enclosure. Sleep deprivation lasted 2–6 h; mice were then returned to their home cages, where old nests had been removed and new nesting material was provided.

### Sleep deprivation through gentle handling

When mice showed signs of quiescence, the experimenter gently stroked them with a soft brush or lightly tapped the cage to prevent sleep. Sleep deprivation lasted either one or four hours and was immediately followed by nest manipulation.

### Acute stress

Prior to the nesting manipulation at Zeitgeber time (ZT) 7.5, test mice underwent 30 minutes of immobilization stress using a well-ventilated perforated 50-ml centrifuge tube.

### Polysomnographic acquisition and analysis

EEG-EMG signals were derived from the surgically implanted electrodes, amplified (Model 3500, A-M systems) and digitized at a rate of 256 Hz (Vital Recorder, Kissei Comtec America). Subsequently, the signal was filtered (EEG, 0-25 Hz; EMG, 25-50 Hz) and spectrally analyzed via fast Fourier transform using SleepSign for Animal software (Kissei Comtec America).

The data were first annotated semiautomatically into 4-s epochs as wake, non-rapid eye movement (NREM) sleep and rapid eye movement (REM) sleep. The scoring was then visually inspected and validated based on the EEG-EMG waveforms and their respective power spectra, with corrections made as necessary. All scoring was done by investigators blind to the experimental manipulation. We defined wakefulness by a low-amplitude mixed-frequency EEG paired with tonic EMG activity exhibiting phasic bursts. NREM sleep was defined by a high-amplitude, low-frequency (0.5-4 Hz) EEG and a markedly diminished EMG activity relative to wakefulness. REM sleep was defined by diminished low-frequency oscillations, a prominent theta rhythm (4-9 Hz), and an absence of EMG activity.

To determine the latency to NREM sleep, we identified the initial NREM sleep episode exceeding 12 seconds. For REM sleep latency, we identified the first REM sleep episode surpassing 8 seconds. For the latency-to-sleep analyses, a maximum latency of 120 min was assigned when mice did not enter the corresponding sleep state during the analysis period. Specifically, this value was assigned to both NREM and REM sleep for one mouse that remained continuously awake in Figure 3E and three mice in Figure 5E; and to REM sleep only for three mice in Figure 3E, two mice in Figure 4B, and one mouse in Figure 5E. For analyses of episode duration, only time windows containing at least one episode of the corresponding arousal state were included. Time windows without wake, NREM sleep, or REM sleep episodes were coded as missing values. We excluded data from three outlier mice in the mean wake episode analysis in Figure 3D and from two outlier mice in Figure 4E, as these animals spent over 85% of the 2-h analysis period continuously awake, resulting in mean episode durations greater than 900 sec. In addition, we excluded power density data from four sessions presented in Figure 2F due to noise in the EEG signal.

**Figure 1.**
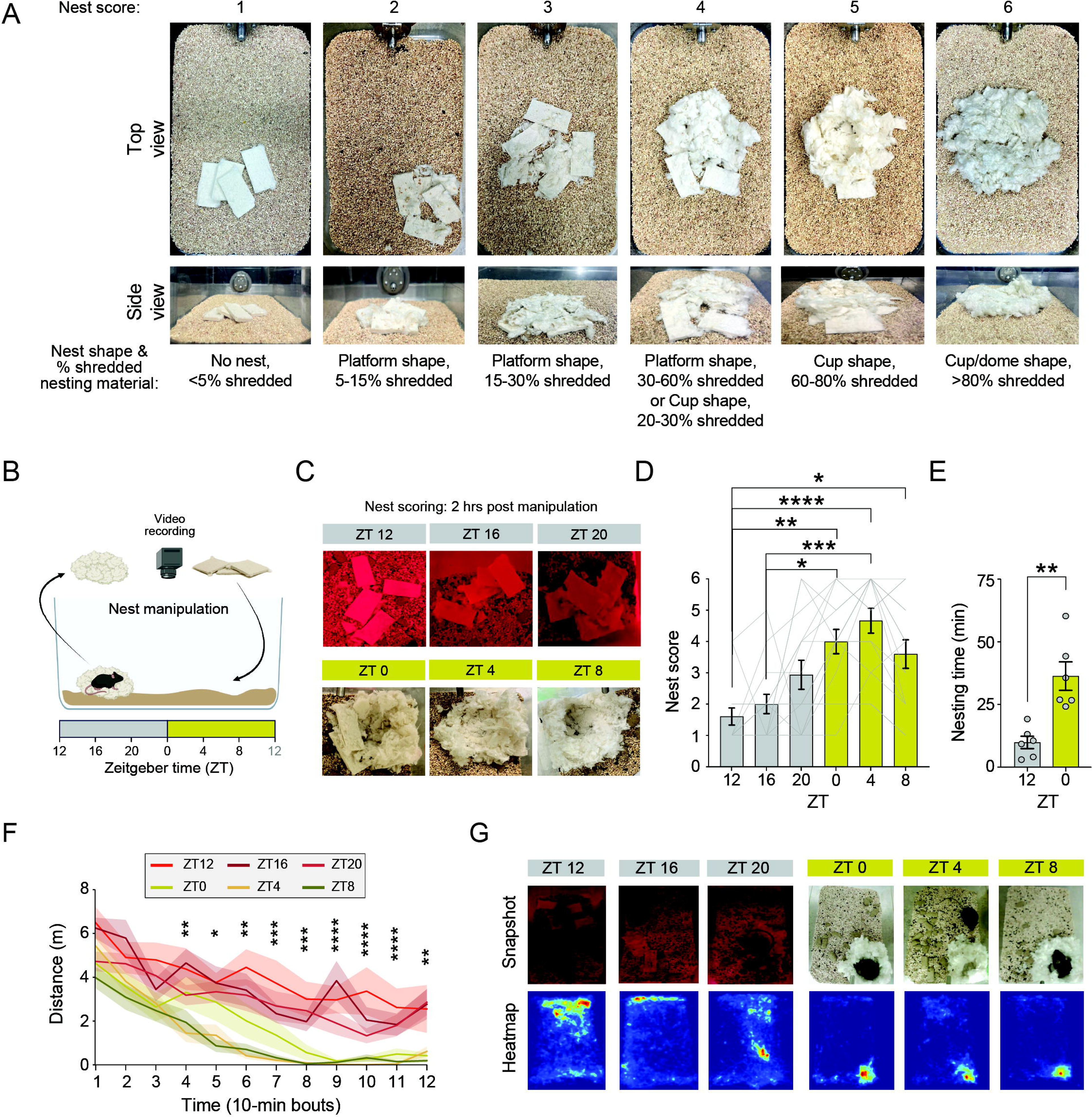
Mice show a strong motivation to build nests throughout the light phase. (**A**) Visual representation of nest scores, with home cage images from top and side views, along with the corresponding nest shape and percentage of nesting material used shown below each image. (**B**) Schematic of the nesting manipulation paradigm. Nests were removed from home cages at different circadian time points (Zeitgeber time [ZT] 12, 16, 20, 0, 4, and 8), and new nesting material (5 g Nestlet) was placed inside each home cage, away from the previous nest location. Nests were scored two hours post-manipulation. Representative images of nests (**C**) and nest scores (**D**) of mice whose nesting material was manipulated at ZT 12, 16, 20, 0, 4, and 8. n = 15 female and male mice. Friedman test followed by Dunn’s multiple comparisons. (**E**) Total time spent nesting during the 2-hour period following nest manipulation at ZT 12 and at ZT 0. n = 6 female and male mice. One-tailed paired t test. (**F**) Distance traveled during the 2-hour post-manipulation period, calculated from bounding box centroids. n = 12 female and male mice. Friedman test followed by False Discovery Rate (FDR)-corrected post hoc Wilcoxon signed-rank tests. (**G**) Randomly selected snapshots (top) and corresponding heatmaps (bottom) of mice following nesting manipulations at ZT 0, 4, 8, 12, 16, and 20. Heatmaps display bounding box centroid locations across the 2-hour post-manipulation period. Data are shown as mean ± SE. ∗, p < 0.05; ∗∗, p < 0.01; ∗∗∗, p < 0.001; ∗∗∗∗, p < 0.0001.

**Figure 2:**
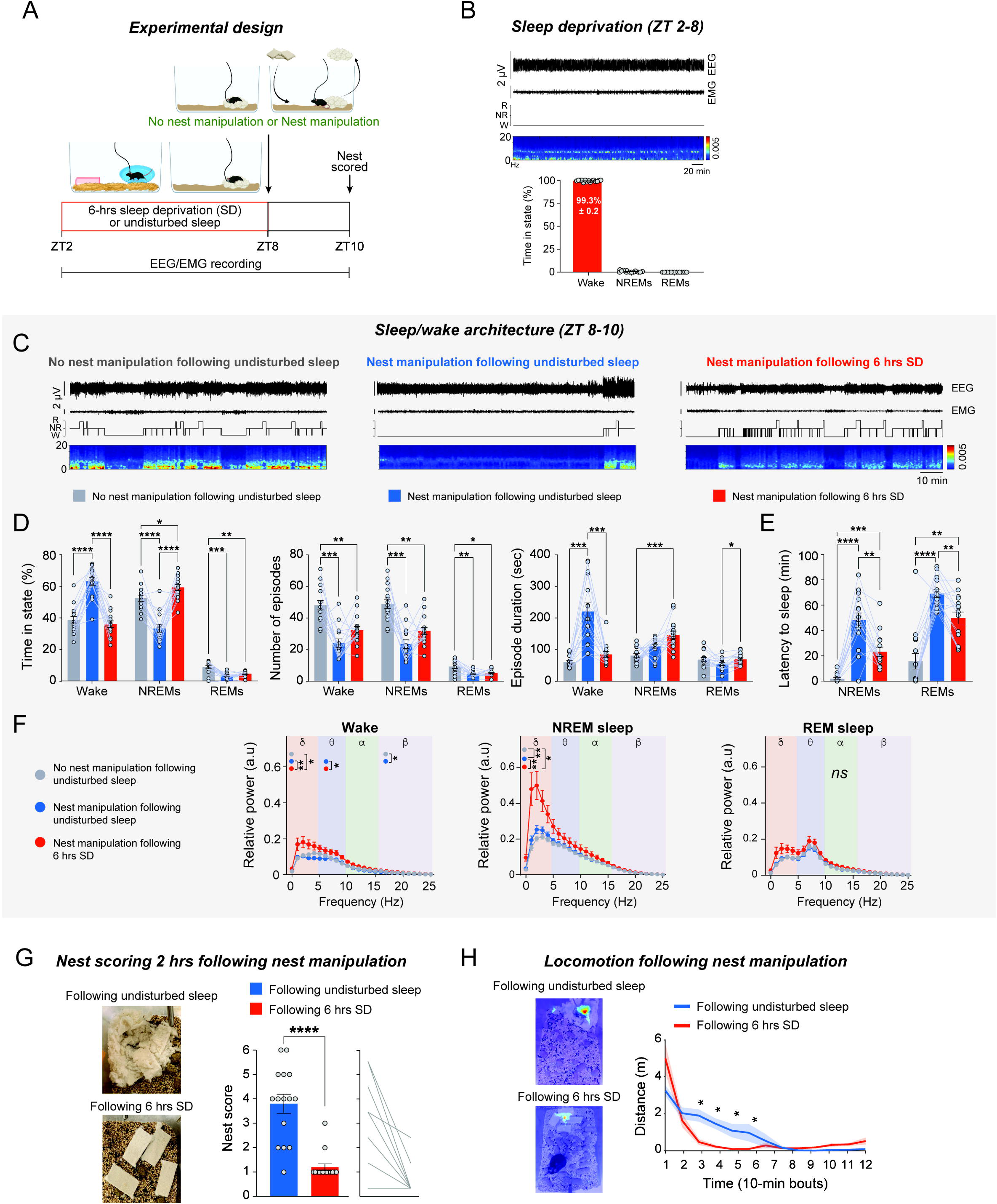
Six hours of sleep deprivation suppresses pre-sleep nest-building behavior. (**A**) Schematic of experimental design. Mice were subjected to six hours of sleep deprivation (SD) via environmental enrichment from Zeitgeber time (ZT) 2–8, followed by nest manipulation at ZT 8. Control mice were either not subjected to SD prior to nest manipulation or were left with their original nests undisturbed from ZT 2–10. EEG, EMG and video data were recorded throughout, and nests were evaluated two hours after manipulation (ZT 10). (**B**) Top: Representative 6-hour EEG and EMG traces, hypnogram, and EEG spectrogram from a mouse subjected to SD. Bottom: Percentage of time spent in wake, NREM sleep, and REM sleep during ZT 2–8 under sleep-deprived conditions. n = 15 female and male mice. (**C**) Representative 2-hour EEG and EMG traces, hypnogram, and EEG spectrogram from ZT 8–10 in mice subjected to the three experimental conditions. (**D**) Sleep/wake architecture during ZT 8–10 in mice subjected to the three experimental conditions. Left: Percentage of time spent in wake, NREM sleep, and REM sleep. Middle: Number of wake, NREM sleep, and REM episodes. Right: Duration of wake, NREM sleep, and REM sleep episodes. n = 15 female and male mice. Two-way repeated-measures (RM) analysis of variance (ANOVA) followed by Tukey’s multiple comparisons tests. (**E**) Latency to NREM and REM sleep. n = 15 female and male mice. A subset of the undisturbed mice (no nest manipulation following undisturbed sleep; n = 9/15) was assigned a latency value of 0, as these animals were already in NREM sleep at the beginning of the analysis period (ZT 8). Two-way RM ANOVA followed by Tukey’s multiple comparisons test. (**F**) Relative EEG power density for wake (left), NREM sleep (middle) and REM sleep (right) episodes during ZT 8–10. Delta = 0–4 Hz; Theta = 5–9 Hz; Alpha = 10–15 Hz; Beta = 16–25 Hz; n = 15 female and male mice. Two-way RM mixed-effects ANOVA followed by Tukey’s multiple comparisons. (**G**) Nest scores 2 hours post-manipulation in mice that were either left undisturbed or subjected to 6 hours of SD. Left: Representative nest images. Right: Mean ± SE with individual mouse data. n = 15 female and male mice. One-tailed Wilcoxon matched-pairs signed-rank test. (**H**) Distance traveled during the 2-hour period following nest manipulation, calculated from bounding box centroids in mice that were either previously undisturbed or subjected to 6 hours of SD. Left: heatmaps of centroid locations during the post-manipulation period. Right: distance data binned into 10-minute intervals. n = 10 female and male mice. Wilcoxon signed-rank tests (FDR-corrected). Data are shown as mean ± SE. ns, p > 0.05; ∗, p < 0.05; ∗∗, p < 0.01; ∗∗∗, p < 0.001; ∗∗∗∗, p < 0.0001.

**Figure 3:**
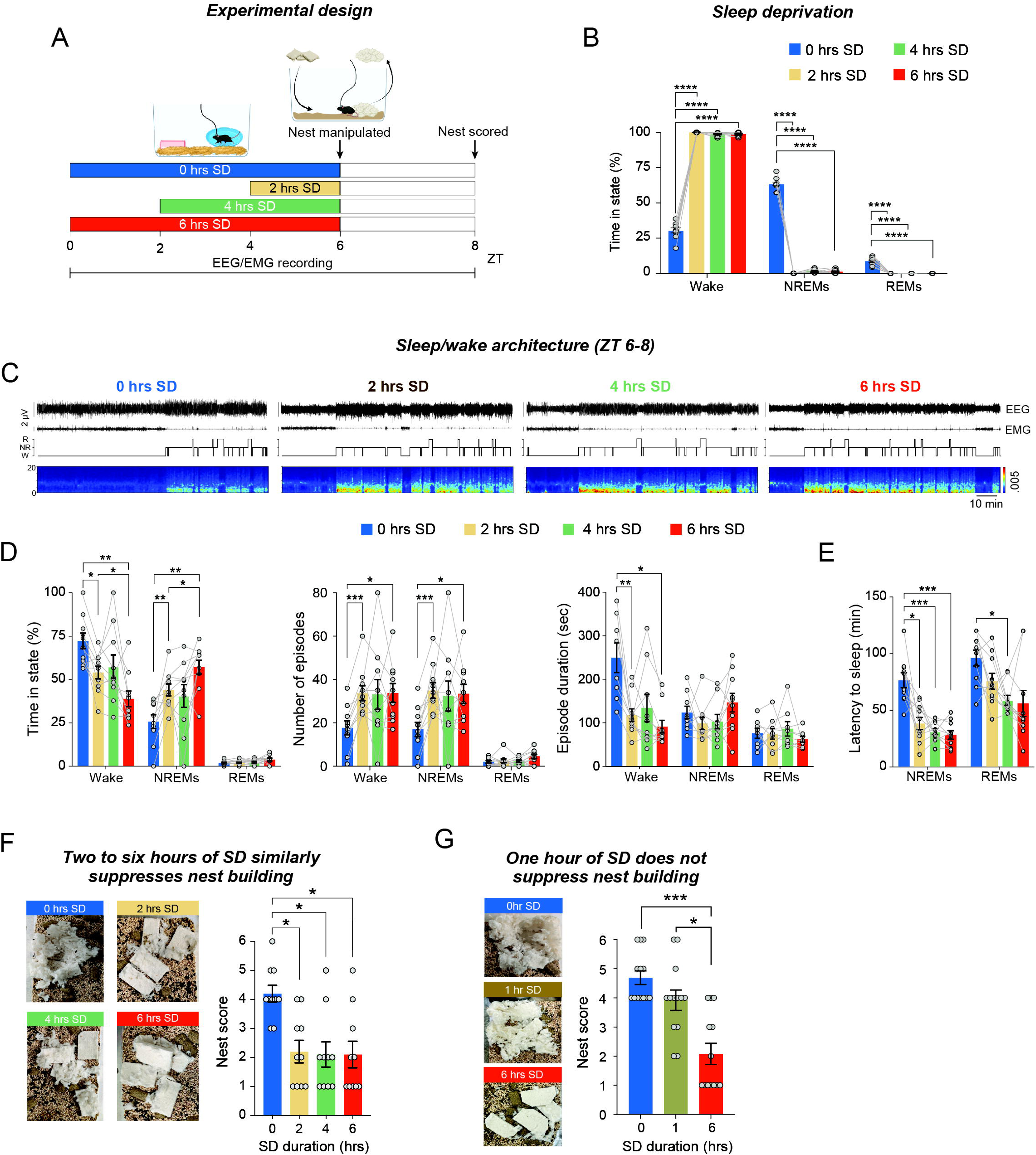
Varying durations of sleep deprivation longer than two hours uniformly suppress pre-sleep nest-building behavior. (**A**) Schematic of experimental design. Mice were subjected to 2, 4, or 6 hours of sleep deprivation (SD) via environmental enrichment, followed by nest manipulation at Zeitgeber time (ZT) 6. Control mice were not sleep-deprived prior to manipulation (‘0 hrs SD’). EEG, EMG and video data were recorded throughout, and nests were evaluated two hours post-manipulation (ZT 8). (**B**) Percentage of time spent in wake, NREM sleep, and REM sleep during the SD procedure. Data for the ‘0 hrs SD’ group correspond to the ZT 0–6 time window. (**C**) Representative 2-hour EEG and EMG traces, hypnogram, and EEG spectrogram from ZT 6–8 in mice subjected to different experimental conditions. (**D**) Sleep/wake architecture during ZT 6–8 following 0, 2, 4, or 6 hours of SD. Left: Percentage of time spent in wake, NREM sleep, and REM sleep. Middle: Number of wake, NREM sleep, and REM sleep episodes. Right: Duration of wake, NREM sleep, and REM sleep episodes. (**E**) Latency to NREM and REM sleep. A maximum latency value of 120 min was assigned for NREM and/or REM sleep in mice that did not enter the corresponding state during the analysis period. (**F**) Left: Representative nest images 2 hours post-manipulation after 0, 2, 4, and 6 hours of SD. Right: Nest scores (mean ± SE with individual mouse data). n = 10 female and male mice. (**G**) Nest scores 2 hours post-manipulation after 0, 1, and 6 hours of SD. Left: Representative nest images. Right: Mean ± SE with individual mouse data. n = 13 female and male mice. For the analyses of percent time in states, number of episodes and latency to sleep, we used two-way RM ANOVA followed by Tukey’s multiple comparisons test. For the analysis of episode duration, we used two-way RM mixed-effects ANOVA, followed by Tukey’s multiple comparisons test. For the analysis of nest scores, we used Kruskal–Wallis test followed by Dunn’s multiple comparisons test. Data are shown as mean ± SE. ns, p > 0.05; ∗, p < 0.05; ∗∗, p < 0.01; ∗∗∗, p < 0.001; ∗∗∗∗, p < 0.0001.

**Figure 4.**
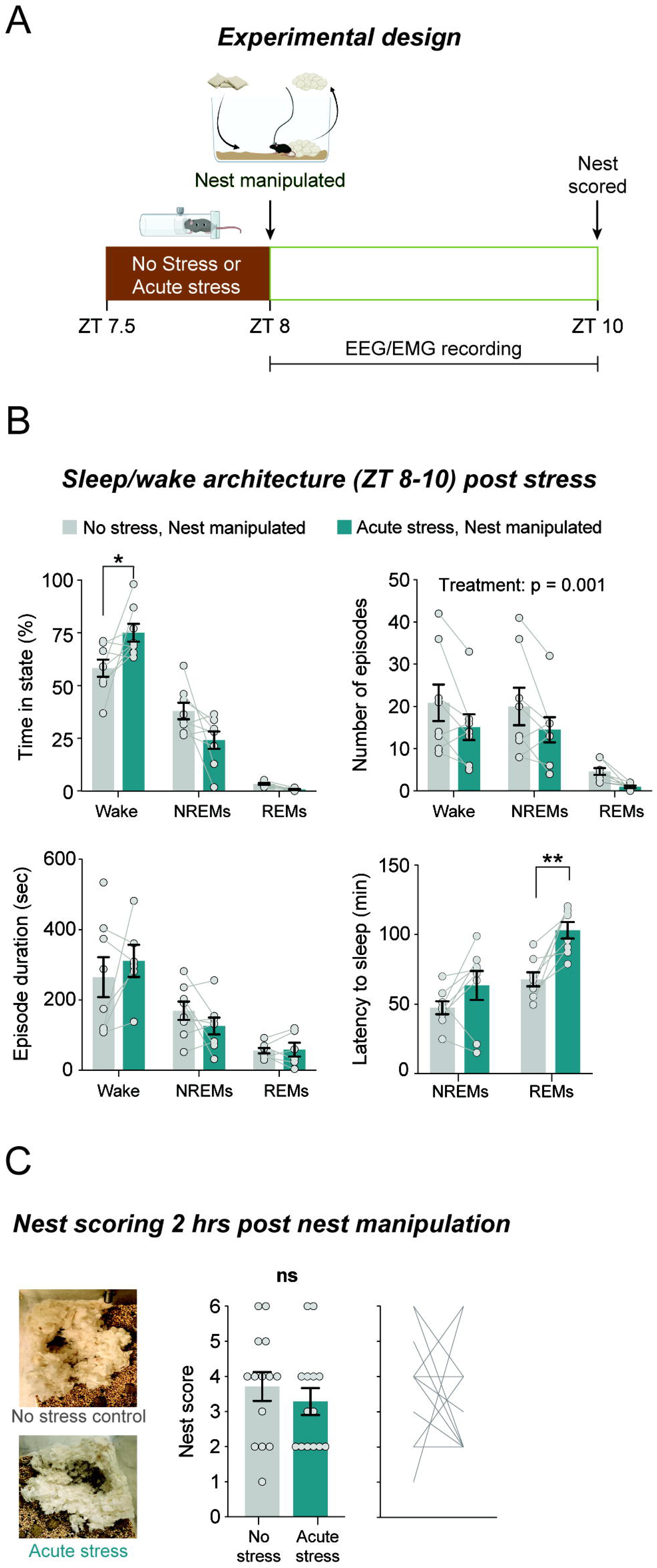
Acute stress does not suppress pre-sleep nest-building behavior. (**A**) Schematic of experimental design. Mice were subjected to 30 minutes of acute restraint stress prior to nest manipulation at Zeitgeber time (ZT) 8 (n = 8 female and male mice). Control mice remained undisturbed before nest manipulation. EEG, EMG, and video data were recorded throughout, and nests were evaluated two hours post-manipulation. (**B**) Sleep/wake architecture during ZT 8–10 in control and stress-exposed mice. Top left: Percentage of time spent in wake, NREM sleep, and REM sleep. Top right: Number of wake, NREM sleep, and REM sleep episodes. Bottom left: Duration of wake, NREM sleep, and REM sleep episodes. Bottom right: Latency to NREM and REM sleep. A maximum latency value of 120 min was assigned for REM sleep in mice that did not enter the state during the analysis period. Data are shown as mean ± SE. For the analyses of percent time in states, number of episodes and latency to sleep, we used two-way RM ANOVA followed by Tukey’s multiple comparisons test. For the analysis of episode duration, we used two-way RM mixed-effects ANOVA, followed by Tukey’s multiple comparisons test. (**C**) Nest scores 2 hours post-manipulation. Left: Representative nest images. Right: Mean ± SE with individual mouse data. One-tailed Wilcoxon matched-pairs signed-rank test. ns, p > 0.05; ∗p < 0.05; ∗∗p < 0.01; ∗∗∗p < 0.001; ∗∗∗∗, p < 0.0001.

**Figure 5:**
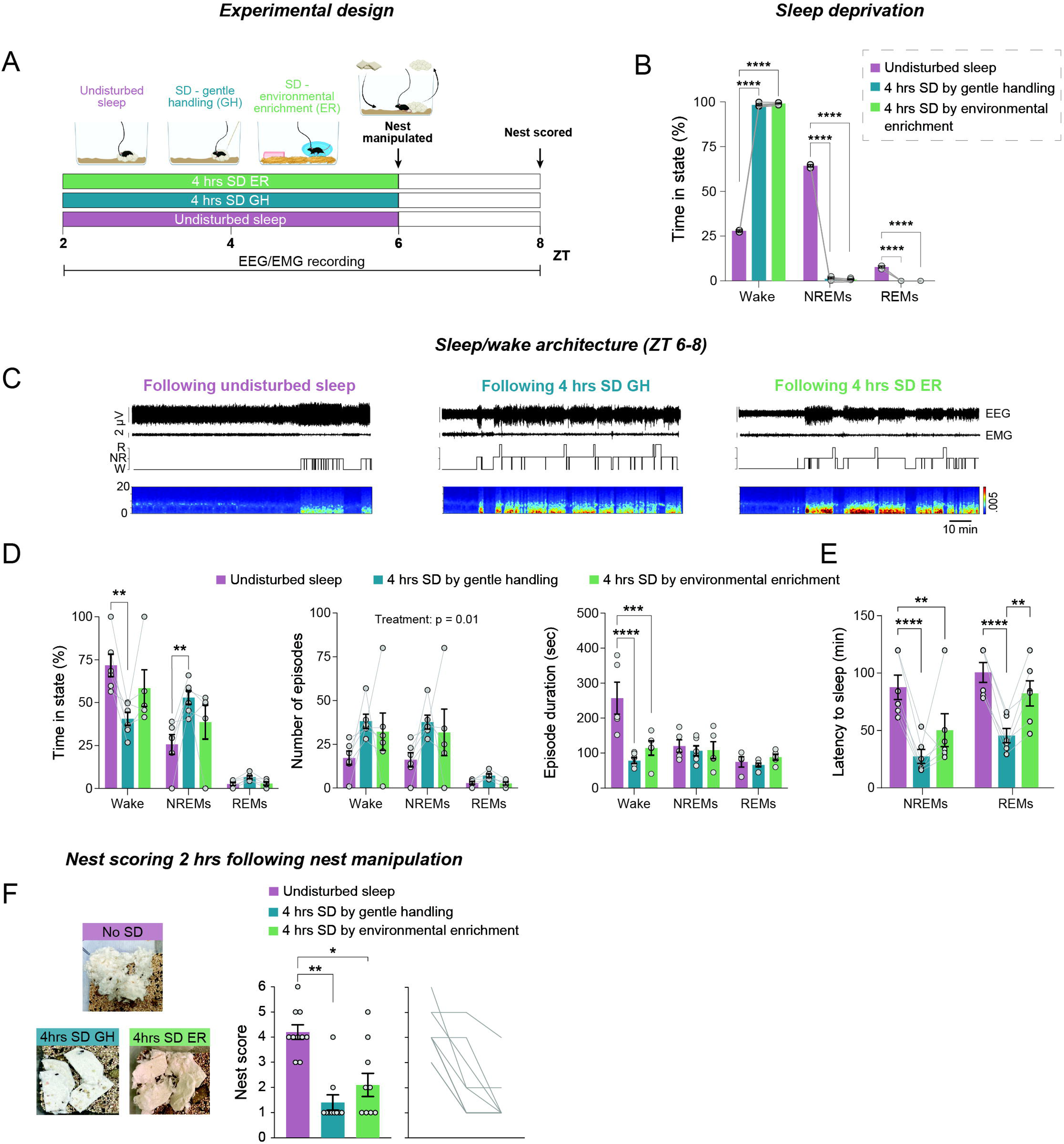
Sleep deprivation by gentle handling suppresses pre-sleep nest-building behavior similarly to sleep. (**A**) Schematic of experimental design. Mice were subjected to 4 hours of sleep deprivation (SD) during Zeitgeber time (ZT) 2–6 via either gentle handling or environmental enrichment, followed by nest manipulation at ZT 6. Control mice were left undisturbed prior to manipulation. EEG, EMG and video data were recorded throughout, and nests were evaluated two hours post-manipulation (ZT 8). (**B**) Percentage of time spent in wake, NREM, and REM sleep during the SD procedure and undisturbed condition. (**C**) Representative 2-hour EEG and EMG traces, hypnogram, and EEG spectrogram from ZT 6–8 in mice subjected to different experimental conditions. (**D**) Sleep/wake architecture during ZT 6–8 in mice subjected to different experimental conditions. Left: Percentage of time spent in wake, NREM sleep, and REM sleep. Middle: Number of wake, NREM sleep, and REM sleep episodes. Right: Duration of wake, NREM sleep, and REM sleep episodes. (**E**) Latency to NREM and REM sleep. A maximum latency value of 120 min was assigned for NREM and/or REM sleep in mice that did not enter the corresponding state during the analysis period. (**F**) Nest scores 2 hours post-manipulation. Left: Representative nest images. Right: Mean ± SE with individual mouse data. n = 6 female and male mice for sleep architecture data, and n = 10 female and male mice for nesting data. We used two-way RM ANOVA, followed by Šídák’s multiple comparisons test for the analysis of percent time in states and number of episodes and by Tukey’s multiple comparisons test for the analysis of sleep latency. We used RM mixed-effects ANOVA followed by Šídák’s multiple comparisons test for the analysis of episode duration, and the Friedman test followed by Dunn’s multiple comparisons for nest scores. ns, p > 0.05; ∗, p < 0.05; ∗∗, p < 0.01; ∗∗∗, p < 0.001; ∗∗∗∗, p < 0.0001.

### Manual analysis of nest-building behavior

We annotated mice nest-building behavior second-by-second using the Behavioral Observation Research Interactive Software (BORIS)^28^ from video data. Nesting was defined for instances where the mouse was engaged in pulling, carrying, fraying, push-digging, sorting or fluffing of nesting material, as in^27^.

### Automatic locomotion analysis

We used the Detect Any Mouse Model (DAMM)^28^ toolbox, which employs instance segmentation to detect and track the location of mice in video recordings, generating bounding boxes that define the position and size of each animal. The original DAMM model weights and configuration files were used in a zero-shot setting^28^. Using custom Python code, we calculated the centroid of each bounding box and computed the mean distance traveled in 10-minute bins. To reduce noise from small posture-related shifts, a minimum displacement threshold of 0.5 cm per frame was applied, and only movements exceeding this threshold were included in the distance calculations. For data presented in Figure 1F, for each time bin, locomotion across ZTs was compared using a Friedman test, followed when significant by False Discovery Rate (FDR)-corrected Wilcoxon signed-rank tests (Supplementary Table 2). For data presented in Figure 2H, for each time bin, locomotion between conditions was compared using a Wilcoxon matched-pairs signed-rank test (FDR-corrected; Supplementary Table 2). We excluded video data from the locomotion analysis for 3/15 mice in Figure 1F and 2/15 mice in Figure 2H due to low visibility during the dark phase.

### Figure and representative video preparation

Data plots were generated using either Prism 9.1 (GraphPad Software) or custom MATLAB (MathWorks, Natick, MA, U.S.A.) or Python scripts and were exported to Adobe Illustrator 2023 (Adobe Creative Cloud) for final figure preparation. Representative videos were created using Adobe Premiere Pro (Adobe Creative Cloud).

### Statistical analysis

Sample sizes were chosen based on prior publications investigating sleep/wake circuitry^6,29,30^. Non-parametric tests were applied for binned or scaled data and for datasets that did not meet normality assumptions. For analyses of variance (ANOVA) with more than two repeated-measures levels, Greenhouse–Geisser correction was applied when appropriate. The Friedman test followed by Dunn’s multiple comparisons test was used for Figures 1D and 5F, and followed by Wilcoxon signed-rank test (FDR-corrected using the Benjamini-Hochberg procedure) was used for Figure 1F. A one-tailed paired t test was used for Figure 1E (because the hypothesis was that nesting time is increased at ZT 0 compared to ZT 12). The Wilcoxon matched-pairs signed-rank test was used for Figures 2G, 2H (FDR-corrected using the Benjamini-Hochberg procedure), and 4C (one-tailed for 2G and 4C, as the hypotheses were that sleep deprivation and stress suppress nest-building). The Kruskal–Wallis test followed by Dunn’s multiple comparisons test was used for Figures 3F and 3G. Two-way repeated-measures (RM) ANOVA was used for Figure S2 (wake). Two-way RM ANOVA followed by Tukey’s multiple comparisons test was used for Figures 2D, 2E, 3B, 3D (percent and number of episodes), 3E, and 5E. Two-way RM ANOVA followed by Šídák’s multiple comparisons test was used for Figures 4B (percent, episode number and latency), 5B, and 5D (percent and number of episodes). Two-way RM mixed-effects ANOVA followed, when appropriate, by Tukey’s multiple comparisons test was used for Figures 2F, 3D (episode duration), 4B (episode duration), S1 and S2 (NREM and REM sleep). Two-way RM mixed-effects ANOVA followed by Šídák’s multiple comparisons test was used for Figure 5D (episode duration). We analyzed data using either Prism 9.1 (GraphPad Software) or custom MATLAB and Python scripts.

## RESULTS

### Mice show a strong motivation to build a nest throughout the light phase

While the need for sleep varies throughout the day^16,17^, it remains unclear whether the motivation to prepare for sleep exhibits similar fluctuations. To explore this, we focused on nest-building behavior, a central pre-sleep activity in mice^6,21,30^. Nest-building can be readily quantified by assessing the shape of the nests and the weight of nesting material shredded within a specific time window (Fig. 1A and ^30,31^). Consequently, the quality of the nest can be used to infer the mice’s motivation to build it.

We subjected singly housed adult mice (n = 7 male and 8 female mice) to nest manipulation by replacing their old nests with new nesting material at various times across the light/dark cycle (ZT 12, 16, 20, 0, 4, and 8). This approach allowed us to evaluate the effects of both the phase of the light/dark cycle and time of day on their motivation to build a nest. We video-recorded the mice and assessed the quality of the nests two hours post-manipulation (Fig. 1B). We found that mice show a daily rhythm in their motivation to build a nest (Fig. 1C**,D**). During the dark phase, specifically at ZT 12 and 16, the mice show reduced motivation to build nests, minimally interacting with the provided nesting material (Fig. 1C**,D**). In contrast, mice demonstrate a consistently strong motivation to build nests across the light phase, with scores peaking at ZT 4 (Fig. 1C), despite the homeostatic drive for sleep being understood to decrease over this period as sleep need diminishes with time spent asleep^16,17^. To verify that our nest scoring scheme accurately represents the time spent nest building, we manually annotated the 2-h-long videos of mice following the nest manipulation conducted at ZT 12 and 0 (n = 3 male and 3 female mice; Fig. 1E). We found that mice spend significantly more time engaged in nest building at ZT 0 compared to ZT 12 (Fig. 1E), as reflected in our nest scoring (Fig. 1D). Together, these findings strongly suggest that mice exhibit a daily oscillation in their motivation to build nests, maintaining consistently high motivation throughout the entire light phase.

Next, using the machine-learning-based mouse tracking system DAMM^28^, we characterized locomotion in our video data during the 2-hour period following nest manipulations conducted at six different time points across the light/dark cycle (n = 6 male and 6 female mice; Fig. 1F). At all ZT times, nest manipulation was followed by robust locomotion lasting for approximately 30 minutes (Fig. 1F **and Supplementary Tables 1 and 2**). Beyond this period, locomotion progressively decreased when manipulation occurred during the light phase, and from 40 minutes post-manipulation onward it differed significantly from the dark-phase conditions (Fig. 1F **and Supplementary Table 2**). Moreover, during the light phase, mice predominantly resided in the final nest location (Fig. 1G; summary data not shown). Together, these findings suggest that nest manipulation induces a transient increase in locomotion regardless of time of day, but during the light phase, this activity subsides once nest-building is completed and mice become quiescent inside their nests.

### Sleep-preparatory nest-building behavior is suppressed by six hours of sleep deprivation

Given that fatigue in humans is known to increase the likelihood of abrupt transitions from wakefulness to sleep (e.g., ^13^)–eliminating the behavioral preparation for sleep–we wondered whether sleep loss would similarly suppress behavioral preparation for sleep in mice. To investigate this, we implanted adult mice (n = 9 male and 6 female mice) with EEG-EMG recording electrodes. After recovery, habituation, and single housing, mice were subjected to six hours of sleep deprivation via environmental enrichment, from ZT 2 to 8 (Fig. 2A). The procedure involved placing each mouse in an enclosure with bedding, water, and a running wheel—but without nesting material or food—and introducing novel objects (e.g., plastic figures) whenever the mouse showed signs of quiescence. We then (at ZT 8) performed nest manipulations, as described above, and assessed the quality of the nests two hours post-manipulation (Fig. 2A). As controls, we used two conditions: six hours of undisturbed sleep followed by nest manipulation, and six hours of undisturbed sleep followed by no manipulation. EEG-EMG and video data were recorded throughout.

We first validated the effectiveness of our sleep deprivation procedure and found that mice subjected to sleep deprivation remained predominantly awake (99.26% ± 0.23% time awake) throughout the entire 6-hour procedure, spending very little time asleep (Fig. 2B).

Next, we characterized the sleep/wake architecture during ZT 8–10, the two hours following nest manipulation for the mice undergoing these manipulations (Fig. 2C**-F**). Mice subjected to nest manipulation after six hours of undisturbed sleep spent significantly more time awake and less time in both NREM and REM sleep compared to mice not subjected to nest manipulation after six hours of undisturbed sleep (Fig. 2D). The increase in wake time was driven by a reduced number of NREM and REM sleep episodes, rather than changes in their mean duration (Fig. 2D). Notably, six hours of sleep deprivation suppressed the effects of nest manipulation on sleep/wake states: mice subjected to nest manipulation after sleep deprivation showed decreased wakefulness and increased NREM sleep compared to those subjected to nest manipulation after undisturbed sleep (Fig. 2C**,D**). While nest manipulation increased the latency to both NREM and REM sleep regardless of prior sleep history, the extent of this delay was reduced in mice subjected to sleep deprivation before the manipulation (Fig. 2E). Together, these results suggest that sleep deprivation diminishes the arousal response typically induced by the need to prepare for sleep.

Six hours of sleep deprivation followed by nest manipulation resulted in a significant increase in delta and theta power during wakefulness compared to the non–sleep-deprived conditions (Fig. 2F). Delta power was also significantly elevated during NREM sleep after six hours of sleep deprivation followed by nest manipulation relative to non–sleep-deprived mice (Fig. 2F). Nest manipulation following undisturbed sleep—and the associated delay in sleep onset—produced a smaller-magnitude yet significant increase in delta power during NREM sleep compared with the undisturbed condition (Fig. 2F). Neither sleep deprivation nor nest manipulation affected EEG power density during REM sleep (Fig. 2F). Taken together, these results are consistent with the well-documented elevation in delta activity during rebound sleep.

We next examined the effects of sleep deprivation on sleep-preparatory nest-building behavior. We found that sleep-deprived mice had significantly lower nest scores two hours post-manipulation compared to mice that underwent nest manipulation at the same circadian time following undisturbed sleep (Fig. 2G **and Supplementary Video 1**). Moreover, mice subjected to nest manipulation after sleep deprivation exhibited a significant reduction in locomotion within 30 minutes of the manipulation, whereas non–sleep-deprived mice showed a comparable decrease only after approximately one hour (n = 9 male and 6 female mice; Fig. 2H). Together, these findings strongly suggest that sleep deprivation suppresses the motivation to build a nest prior to sleep initiation, reducing locomotion and directly promoting sleep.

### Even brief episodes of sleep deprivation, as little as two hours, suppress sleep-preparatory nest-building behavior

We next wondered how different durations of sleep deprivation affect pre-sleep nest-building behavior. To address this, we subjected singly housed mice (n = 5 male and 5 female mice) implanted with EEG-EMG recording electrodes to nest manipulations immediately following 0, 2, 4, or 6 hours of sleep deprivation via environmental enrichment (Fig. 3A). All sleep deprivation procedures ended at ZT 6 to control for potential circadian effects on nest-building behavior. EEG-EMG and video data were recorded simultaneously.

We first validated the effectiveness of our sleep deprivation procedure and found that mice subjected to 2, 4, and 6 hours of sleep deprivation remained predominantly awake (99.2□±□0.22% time awake) throughout the entire procedure, spending very little time asleep compared to mice in the undisturbed sleep condition (Fig. 3B). Next, we characterized the sleep/wake architecture during ZT 6–8, the two-hour period following nest manipulation (Fig. 3C**-E** **and Supp Fig. S1**). We found that the duration of sleep deprivation affected sleep/wake architecture: 2 hours of deprivation resulted in more sleep than 0 hours, but less than 6 hours (Fig. 3D). NREM sleep latency following 2, 4, and 6 hours of deprivation was similarly shorter than after 0 hours of sleep deprivation, whereas REM sleep latency was significantly reduced only following 4 hours of sleep deprivation (Fig. 3E). Notably, EEG power density during wake, NREM sleep, and REM sleep was similar after nest manipulation, irrespective of the preceding duration of sleep deprivation (**Supp Fig. S1**). Next, we examined nest quality following the different durations of sleep deprivation. Surprisingly, we found a similar suppression of nest-building behavior following 2, 4, and 6 hours of sleep deprivation, with nest scores significantly reduced compared to those observed following undisturbed sleep (Fig. 3F **and Supplementary Video 2**). Together, these findings suggest that while sleep debt proportionally modulates sleep/wake architecture, its accumulation uniformly suppresses the motivation to prepare for sleep.

Finally, we wondered how periods of sleep deprivation shorter than two hours would affect nest-building behavior. We repeated the experiment, subjecting a new cohort of mice to nest manipulation after 0, 1, or 6 hours of sleep deprivation (n = 8 male and 5 female mice). Notably, one hour of sleep deprivation did not significantly diminish nest-building, as mice built nests of comparable quality to those of control mice (0 hours of SD) and achieved significantly higher scores than those observed after 6 hours of sleep deprivation (Fig. 3G). These findings suggest that a tipping point in nest-building motivation is reached within 1–2 hours of sleep loss.

### Acute stress does not account for the reduction in nest-building following sleep deprivation

Since sleep deprivation not only suppresses sleep but also alters homeostasis and induces stress, we wondered whether the reduced motivation to build a nest following sleep deprivation might be solely a consequence of stress exposure rather than sleep loss. To test this hypothesis, we subjected singly housed mice (n = 2 male and 6 female mice) implanted with EEG-EMG recording electrodes to 30 minutes of acute restraint stress immediately prior to nest manipulation at ZT 7.5 (Fig. 4A). As a control condition, mice underwent nest manipulation following undisturbed sleep (no-stress condition). EEG-EMG and video data were recorded throughout, and nest quality was assessed two hours post-manipulation.

We first examined sleep/wake architecture during the 2-hour period following nest manipulation. Mice exposed to acute restraint stress prior to the nest manipulation spent more time awake compared to non-stressed controls (Fig. 4B). Stress exposure reduced the number of arousal state episodes, without significantly affecting their mean duration (Fig. 4B). REM sleep latency was significantly increased following acute stress, with a non-significant trend toward increased NREM sleep latency (Fig. 4B). These findings suggest that acute stress elicits an additional arousal response beyond that produced by the nest manipulation itself (**Figs. 2-3**).

We next assessed the effects of acute restraint stress on sleep-preparatory nest-building behavior. Stress exposure did not alter this behavior, as mice exposed to acute restraint stress built nests of comparable quality to those of control mice and achieved similarly high scores (Fig. 4C). Taken together, these findings suggest that sleep loss—rather than acute stress—suppresses the motivation to build a nest prior to sleep.

### Environmental enrichment does not account for the suppression of nest-building behavior following sleep deprivation

Lastly, we wondered whether the enriched experience during sleep deprivation, rather than sleep loss itself, could account for the suppression of nest-building behavior. To test this, we subjected singly housed mice implanted with EEG-EMG recording electrodes (n = 4 male and 2 female mice) to 4 hours of sleep deprivation via gentle handling, from ZT 2 to 6, followed by nest manipulation at ZT 6 (as described above) (Fig. 5A). The sleep deprivation via gentle handling was conducted in the home cage and did not involve the introduction of any salient stimuli, aside from the occasional brush used to gently arouse the mouse. As controls, prior to nest manipulation mice were either allowed to sleep undisturbed or were sleep-deprived for the same duration using environmental enrichment. EEG-EMG and video data were recorded throughout, and nest quality was assessed two hours post-manipulation (Fig. 5A).

We first validated the effectiveness of our sleep deprivation procedures and found that mice subjected to 4 hours of sleep deprivation via either gentle handling or environmental enrichment, remained predominantly awake (98.74□±□0.3% time awake) throughout the procedure, spending very little time asleep compared to mice in the undisturbed sleep condition (Fig. 5B). Next, we characterized sleep/wake architecture during ZT 6–8 (Fig. 5C**-E** **and Supp Fig. S2**) and found that wakefulness was reduced and NREM sleep was increased following sleep deprivation–particularly when conducted via gentle handling–compared to nest manipulation after undisturbed sleep (Fig. 5D). EEG power density was not significantly modulated by the method of sleep deprivation (**Supp Fig. S2**). Sleep deprivation by either method reduced the latency to sleep following nest manipulation relative to the undisturbed sleep condition (Fig. 5E). Finally, we examined nest quality following the different methods of sleep deprivation (n = 5 male and 5 female mice). We found that regardless of the deprivation method, 4 hours of sleep deprivation suppressed sleep-preparatory nest-building behavior, with nest scores significantly reduced compared to those observed following undisturbed sleep (Fig. 5F **and Supplementary Video 3**). This finding strongly suggests that sleep loss–rather than the enriched experience during the deprivation procedure–is responsible for suppressing the motivation to build a nest prior to sleep.

## DISCUSSION

Our study reveals that the motivation to engage in pre-sleep nest-building behavior in mice follows a daily rhythm. Mice consistently show high nest-building motivation during the light phase–despite variations in homeostatic sleep drive–and markedly reduced motivation during the dark phase. By systematically excluding alternative explanations, we demonstrate that sleep loss overrides nest-building behavior prior to sleep initiation, preventing the organism from organizing a safe sleeping environment and instead imposing sleep directly. Furthermore, we show that varying amounts of sleep loss, from two to six hours, uniformly suppress nest-building, suggesting that preparatory behaviors are actively regulated by mechanisms distinct from those governing the sleep process itself. Notably, our findings further suggest that within one to two hours of sleep loss, a tipping point is reached that subsequently diminishes the capacity to prepare for sleep. Our work significantly advances current understanding of the processes that regulate the transition from wakefulness to sleep. A deeper understanding of the process of falling asleep is important—not only because abrupt and uncontrolled transitions into sleep can have serious consequences for individuals and society, but also because difficulty initiating sleep is a core symptom of insomnia^24^ and a strong predictor of cognitive decline later in life^23,24^.

By examining the motivation to build a nest at six time points across the 24-hour light/dark cycle, we show that mice exhibit a clear daily rhythm in their drive to engage in pre-sleep nest-building behavior. This rhythm appears to be more closely linked to the timing of sleep than to the homeostatic sleep need, as nest-building motivation remains high throughout the light phase—even as sleep pressure naturally dissipates^16,17^. Furthermore, our sleep deprivation experiments, which show a similar suppression of nest-building motivation following two, four, and six hours of deprivation, suggest that the sleep pressure induced by deprivation may differ qualitatively from that which accumulates during typical wakefulness in the active phase. How the neuronal substrates underlying homeostatic and circadian sleep drives interact with circuits mediating pre-sleep motivation remains a key open question.

While much effort over recent decades has focused on elucidating the roles of circadian and homeostatic processes in regulating sleep timing, duration, and depth, their influence on pre-sleep behaviors–and the neurobiological mechanisms that govern these behaviors–remains largely unexplored. This gap is striking, given that preparatory behaviors not only consistently precede sleep initiation but are also tightly and temporally aligned with it, suggesting that they are regulated by closely interacting physiological systems.

Several neuronal populations have been implicated in the regulation of pre-sleep nest-building behavior and sleep initiation, including dopaminergic neurons of the ventral tegmental area (VTA)^30^, somatostatin (SST)-expressing neurons in the medial prefrontal cortex (mPFC)^32^, and a glutamatergic subpopulation in the lateral hypothalamus (LH)^6^. VTA dopaminergic neurons regulate both sleep/wake states and behavioral preparation for sleep; reduced activity in these neurons permits pre-sleep behaviors such as nest-building, whereas their activation suppresses them^30^. In contrast, activation of SST-expressing mPFC neurons promotes both nest-building and sleep^32^. We have recently identified a glutamatergic neuronal subpopulation in the LH that regulates the motivation to engage in pre-sleep nest-building^6^. Chemogenetic activation of this ensemble promotes nest-building behavior, while its inhibition suppresses it^6^.

Notably, suppressing the activity of these neurons also promotes sleep–but only when mice are confronted with the need to prepare a space in which to sleep^6^. In this context, activation of the ensemble promotes wakefulness, while inhibition leads to a rapid transition into sleep^6^. These findings suggest that this LH ensemble supports arousal and goal-directed, sleep-related behaviors specifically in the face of rising sleep pressure. Notably, our current findings on the suppression of nest-building following sleep loss closely resemble the effects observed with chemogenetic inhibition of this LH subpopulation^6^. Whether this behavioral suppression reflects a physiological inhibition of the same neuronal ensemble during sleep deprivation remains to be determined.

As sleep is a vulnerable state associated with reduced thermoregulation and increased exposure to predators and environmental threats, it is advantageous to restrict its occurrence to safe and ecologically appropriate settings. Preceding sleep with behaviors involving locating and preparing a sleeping area–that warrant that sleep is initiated under the proper conditions, such as within a nest–may represent an efficient evolutionary strategy. It is therefore striking that animals–including humans–can sometimes lose the ability to properly prepare for sleep, instead falling asleep abruptly even in dangerous situations. This phenomenon suggests that once sleep pressure rises beyond a critical level, the organism can no longer mount the appropriate arousal response needed to prepare for sleep or resist the drive to fall asleep, regardless of environmental context. Abrupt transitions into sleep likely reflect a fundamental physiological necessity; yet the neural mechanisms responsible for overriding sleep preparatory behaviors remain to be elucidated.

The *de-arousal model of sleep initiation^7^* proposes that the expression of repetitive pre-sleep behaviors, such as grooming and nest-building, reduces vigilance toward the external environment, thereby lowering the tone of wake-promoting neuromodulators. This reduction in neuromodulatory tone leads to a brain-wide shift in excitability, functional connectivity, and information flow, ultimately imposing sleep upon the organism^7^. Our current findings suggest that sleep loss bypasses this usual preparatory step and triggers sleep directly. Future studies could critically test this hypothesis.

Here, we provide the first in-depth characterization of the relationship between sleep deprivation and pre-sleep behavior in mice, laying the groundwork for future mechanistic studies. Given that difficulty falling asleep and undesired transitions into sleep are widespread concerns in multiple sleep and neurodegenerative disorders^34^, as well as during aging–and are predictors of cognitive decline later in life^23,24^–a deeper understanding of the neural mechanisms that regulate the natural transition into sleep, and how these processes are altered under conditions of sleep loss, is of significant interest for the development of more effective therapeutic strategies.

## ACKNOWLEDGMENTS

We thank Autumn Cain and Chelsea Markunas for assistance with data collection and scoring. This work was supported by the National Institute of Neurological Disorders and Stroke (R01 NS131821 and R01 NS129874, A.E.-R.). Cage illustrations appearing in Figures 1B, 2A, 3A, 4A, and 5A were created using BioRender.

## DISCLOSURE STATEMENT

Financial Disclosure: None.

Non-Financial Disclosure: None.

A previous version of this manuscript was uploaded to BioRxiv.

## DATA AND CODE AVAILABILITY

All code will be available via GitHub. All data will be provided upon reasonable request. Any additional information required to reanalyze the data reported in this paper is available from the lead contact upon request.

