## Supplementary Figure Legend for "Sleep loss overrides sleep-preparatory behavior to promote sleep"

**Supplementary Figure S1**: **EEG power density during ZT 6–8 following nest manipulation after 0, 2, 4, and 6 hours of sleep deprivation.** Left: Wake episodes, Middle: NREM sleep episodes, Right: REM sleep episodes. Delta = 0–4 Hz; Theta = 5–9 Hz; Alpha = 10–15 Hz; Beta = 16–25 Hz; n = 10 female and male mice. Data are shown as mean ± SE. Two-way RM mixed-effects ANOVA. ns, p > 0.05.

**Supplementary Figure S2**: **EEG power density during ZT 8–10 following nest manipulation after 4 hours of sleep deprivation by gentle handling and by environmental enrichment.** Left: Wake episodes, Middle: NREM sleep episodes, Right: REM sleep episodes. Delta = 0–4 Hz; Theta = 5–9 Hz; Alpha = 10–15 Hz; Beta = 16–25 Hz; n = 6 female and male mice. Data are shown as mean ± SE. Two-way RM mixed-effects ANOVA. ns, p > 0.05.

**Supplementary Table 1.** Summary of statistical analyses for Figures 1–5.

**Supplementary Table 2.** Summary of locomotion statistics for Figures 1F (top) and 2H (bottom). Top: Friedman χ² statistics and FDR-corrected Wilcoxon signed-rank post hoc p values for locomotion across ZTs, computed separately for each time bin (n = 12 mice). Bottom: Wilcoxon matched-pairs signed-rank statistics (W) and FDR-corrected p values for locomotion following nest manipulation after 0 and 6 h of sleep deprivation, computed separately for each time bin (n = 10 mice).
