## Supplementary figures and images for "Sleep loss overrides sleep-preparatory behavior to promote sleep"

### Figure S1

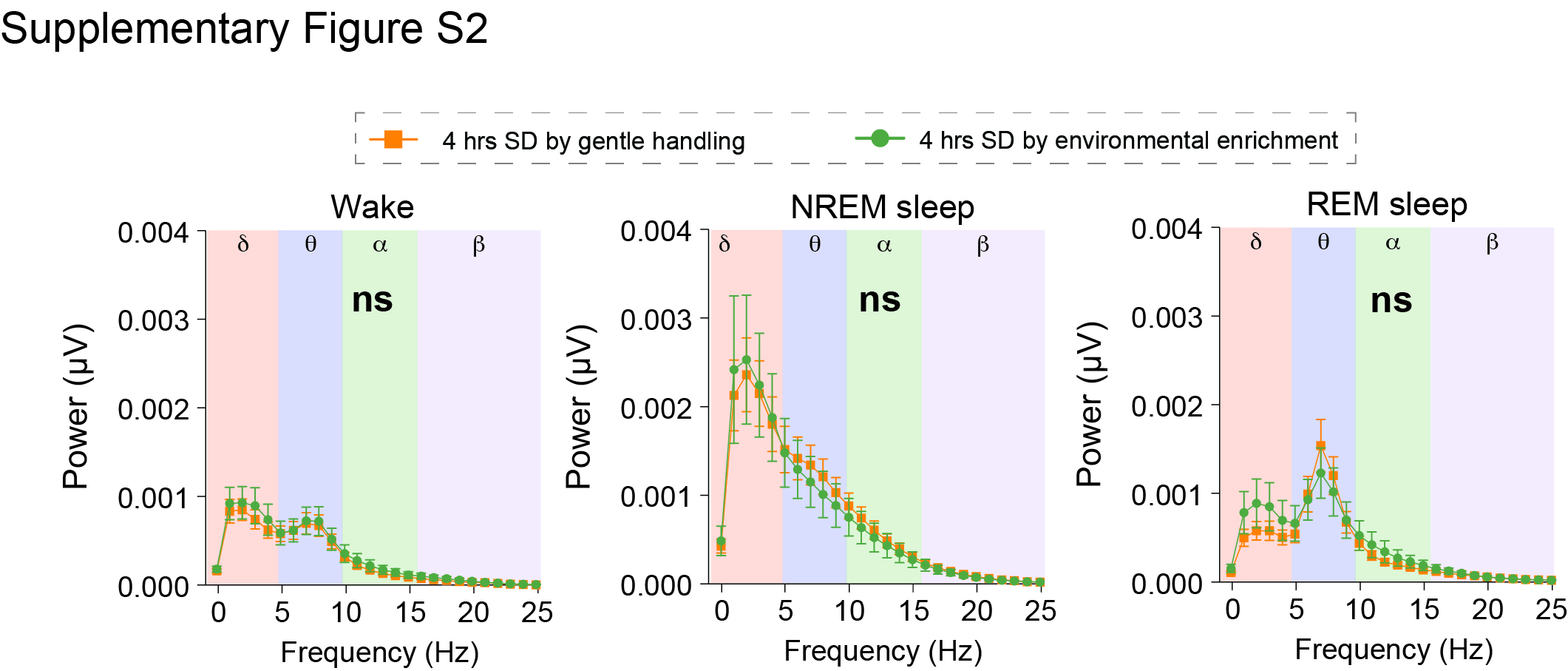

### Figure S2

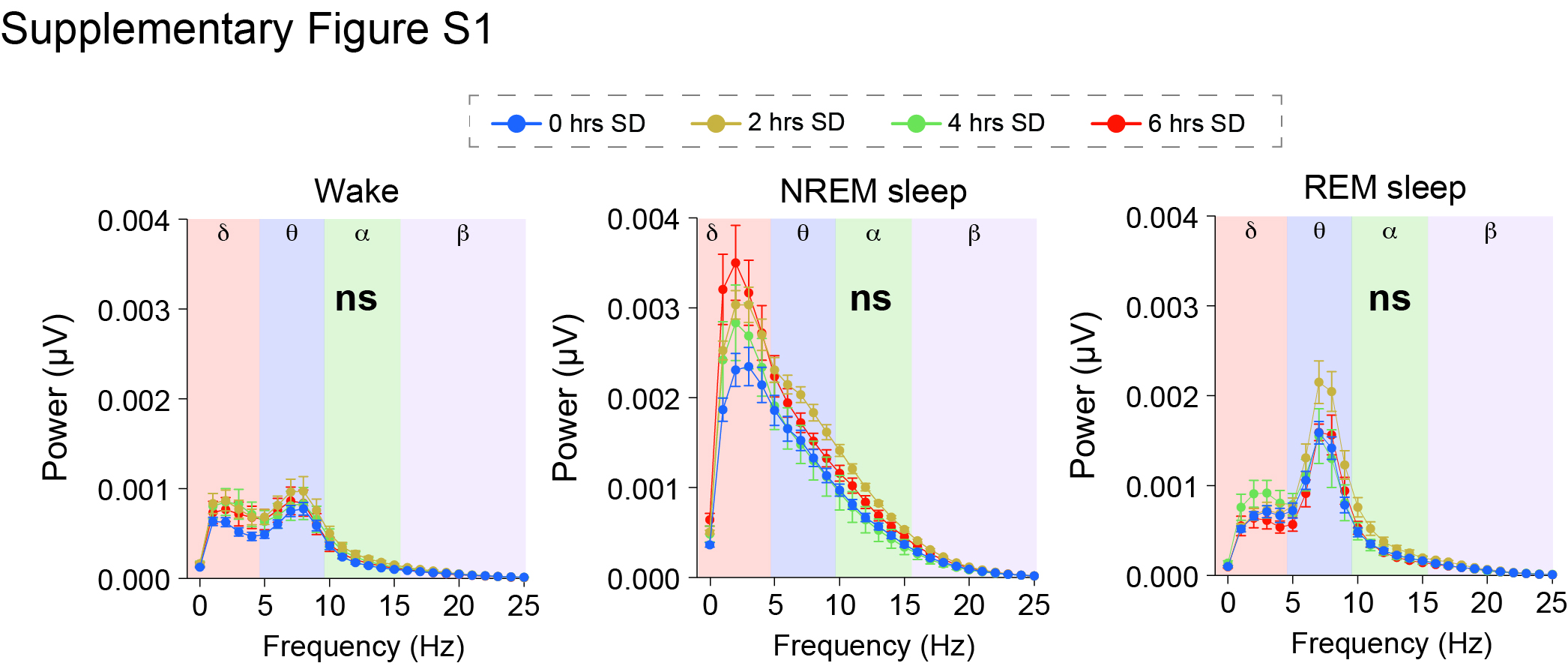
